# Cell-type specific prediction of RNA stability from RNA-protein interactions

**DOI:** 10.1101/2024.11.19.624283

**Authors:** Sepideh Saran, Svetlana Lebedeva, Antje Hirsekorn, Uwe Ohler

**Affiliations:** The Berlin Institute for Medical Systems Biology, Max Delbrück Center for Molecular Medicine, Germany; Department of Biology, Humboldt Universität zu Berlin, Germany; Department of Computer Science, Humboldt Universität zu Berlin, Germany; Department of Electrical Engineering and Computer Science, Technical University of Berlin, Germany

**Keywords:** RNA-binding proteins, RNA stability, translation, machine learning, interpretability

## Abstract

RNA-binding proteins (RBPs) are important contributors to post-transcriptional regulatory processes. The combinatorial action of expressed RBPs and non-coding factors bound to the same transcript determines post-transcriptional properties of the mRNA in a context-dependent manner. To gain a better understanding of RNA stability and translational activity across different conditions, we have compiled and analyzed a set of ribosome profiling datasets for four human cell lines and used existing and newly generated metabolic labeling data to determine matching RNA degradation rates. We then used machine learning methods to predict RNA degradation rate and translation level from RBP binding information, which comprised existing in vivo binding datasets and computationally predicted binding sites. Utilizing this new RNA stability resource, we predicted RNA degradation rate and translation level from RBP binding alone. In vivo binding sites had higher importance for prediction compared to computationally predicted binding sites, likely due to confounding effects. We further explored the feature importance of different RBPs for stability prediction in the context of differential stability conferred by 3’UTR isoforms. Taken together, an RNA stability machine learning model trained on one context successfully generalizes but is impacted by the availability and reliability of current data.

## 1. Introduction

Gene expression is regulated at multiple levels to ensure the correct temporal and cell-typespecific development and homeostasis of an organism. Post-transcriptional mechanisms, such as those regulating RNA processing, splicing, export, localization, translation, and degradation of the RNA, play an important role in ensuring that the transcriptome is buffered against transcriptional noise [21] and can be quickly remodeled if needed [44]. In addition to non-coding regulatory RNAs, post-transcriptional steps are regulated by RNA-binding proteins (RBPs), which recognize short sequence and/or structure motifs on the RNA and form an RBP code that determines the RNA fate. RBPs interact with effector proteins, assembling a complex with a specific action on the RNA. Competition between RBPs for binding sites and auto-regulatory feedback loops contribute to the gene regulatory landscape [43],[37].

Global RNA stability is a combination of separate nuclear [40] and cytoplasmic RNA decay processes, with cytoplasmic RNA stability strongly influenced by deadenylation rate [15] as well as translation speed and codon composition [14]. There are two main strategies to measure RNA degradation rates in mammalian cells: either measuring leftover RNA at different time points after transcription inhibition [27], or using metabolic labeling with nucleotide analogs and monitoring their incorporation over time, as newly synthesized RNA replaces the degraded old RNA [10], [22]. Both methods have been used in multiple studies measuring RNA stability in human and mouse cell lines, and they generally agree with each other [30].

To determine RBP binding sites, methods that use UV light-induced covalent cross-linking and immunoprecipitation of the RBP coupled with sequencing of the associated RNA (Cross-Linking and Immunoprecipitation sequencing, CLIP-seq) are prevalent [20]. RBP binding sites are called as clusters of accumulated reads, with protocol-specific diagnostic events such as mutations that indicate the cross-linked nucleotide allowing CLIP to have single-nucleotide resolution. With the accumulation of hundreds of CLIP datasets, accompanied by functional data of RBP knockdown experiments, RBP binding preferences and regulatory grammar have been pursued in a systematic way [36, 46].

Challenges associated with CLIP experimental data, include a substantial number of false negatives (due to high dependence on transcript abundance) and false positives (due to shortness of reads and contamination with abundant RNA species). With the rise in using deep learning approaches for genomics applications, convolutional neural networks (CNNs) have been trained on CLIP data to learn RBP binding preferences from RNA sequences [18]. This approach helps filter false negatives present in experimental datasets and to be able to predict binding events in a different cell type in which the experimental data is unavailable.

Prior models have predicted RNA stability from influencing factors such as the RNA sequence and RBP binding events. Yeast RNA half-lives lie within minutes and can be predicted well from sequence alone, with codon composition explaining 55% half-life variability and 3’UTR sequences another 5% [8]. Mammalian RNAs have typical half-lives of several hours, and prediction accuracy is lower. A recent deep learning approach complemented RBP features with codon composition, exon junctions, and 5’UTR features to improve the prediction performance [1].

Most existing datasets for RNA stability report gene-level half-lives averaged across isoforms. However, the expectation is that stability levels of the same gene are regulated by different isoforms [19]. Measuring both isoform-specific stability and RBP binding is not straightforward due to the limited ability of short sequencing reads to resolve isoform composition, and while long-read sequencing techniques show promise in this regard [33, 3], such datasets are very rare. Since many machine learning (ML) methods, especially deep learning models, require large datasets in order to achieve acceptable predictive performance, many studies attempt to aggregate stability datasets across cell lines and even species. This approach, however successful in identifying general patterns common among different data sources, neglects differential regulation across conditions.

In this work, we aim to dissect RNA stability determinants conferred by RBPs. Unlike prior work, we focus on finding differential factors across isoforms and cell lines. To this end, we generated novel stability datasets for several cell lines. We then trained ML models to predict stability from RBP binding information and use ML interpretability methods on the models to identify the most influential RBPs. We explore the limits of existing datasets and the extent of their usability for this task. We employ predicted RBP binding sites to identify differential stability features across isoforms of the same gene. We evaluate our approach by using models trained on one cell line to predict stability on unseen data from other cell lines as well as massively parallel reporter experiments. We conclude that there is a robust component in RNA stability that can be interpreted with the current state of existing data and methods at cell line- and isoform-level.

## 2. Results

### 2.1. Establishing feature maps from ribosome profiling and RNA degradation data

The availability of in vivo target site information for hundreds of RNA-binding proteins opens a possibility to use machine learning to uncover possibly multiple roles of RNA binding proteins on distinct aspects of RNA metabolism. We set out to predict mature RNA degradation rate, translation level, and translation efficiency from RBP binding data in human cells. To this end, we implemented a framework with two main steps: the first part uses DNNs to calculate RBP binding sites for each transcript, the second part defines downstream tasks to use the binding information to learn a regressor for predicting degradation or translation rate. We leverage both in-vivo RBP binding data from CLIP experiments, here referred to as raw CLIP data, and in-silico binding predictions. Our raw CLIP data comprises the PAR-CLIP compendium of RBPs profiled in HEK293 cells [36], as well as the ENCODE eCLIP datasets in K562 and HepG2 cells [46]. These CLIP datasets had recently been used to train DeepRiPe, a convolutional neural network classifier that predicts RBP binding sites along an input RNA sequence [18, 23].

The workflow is summarized in Figure 1. In the first step, feature maps are computed. To this end, each sequence was divided into non-overlapping bins of 150nt or 200nt to be consistent with DeepRiPe’s input size. We then assigned binary binding labels based on raw CLIP data: For each transcript, we assigned one to an RBP class if the transcript contains one or more bins with annotated binding sites binding sites, and zero otherwise. To derive the in-silico features, we ran inference on existing pre-trained DeepRiPe models [18] to predict RBP binding for each sequence bin. We used the maximal predicted score among all bins per transcript or 3’UTR, to aggregate predictions across the bins back to isoform-level.

**Figure 1:**
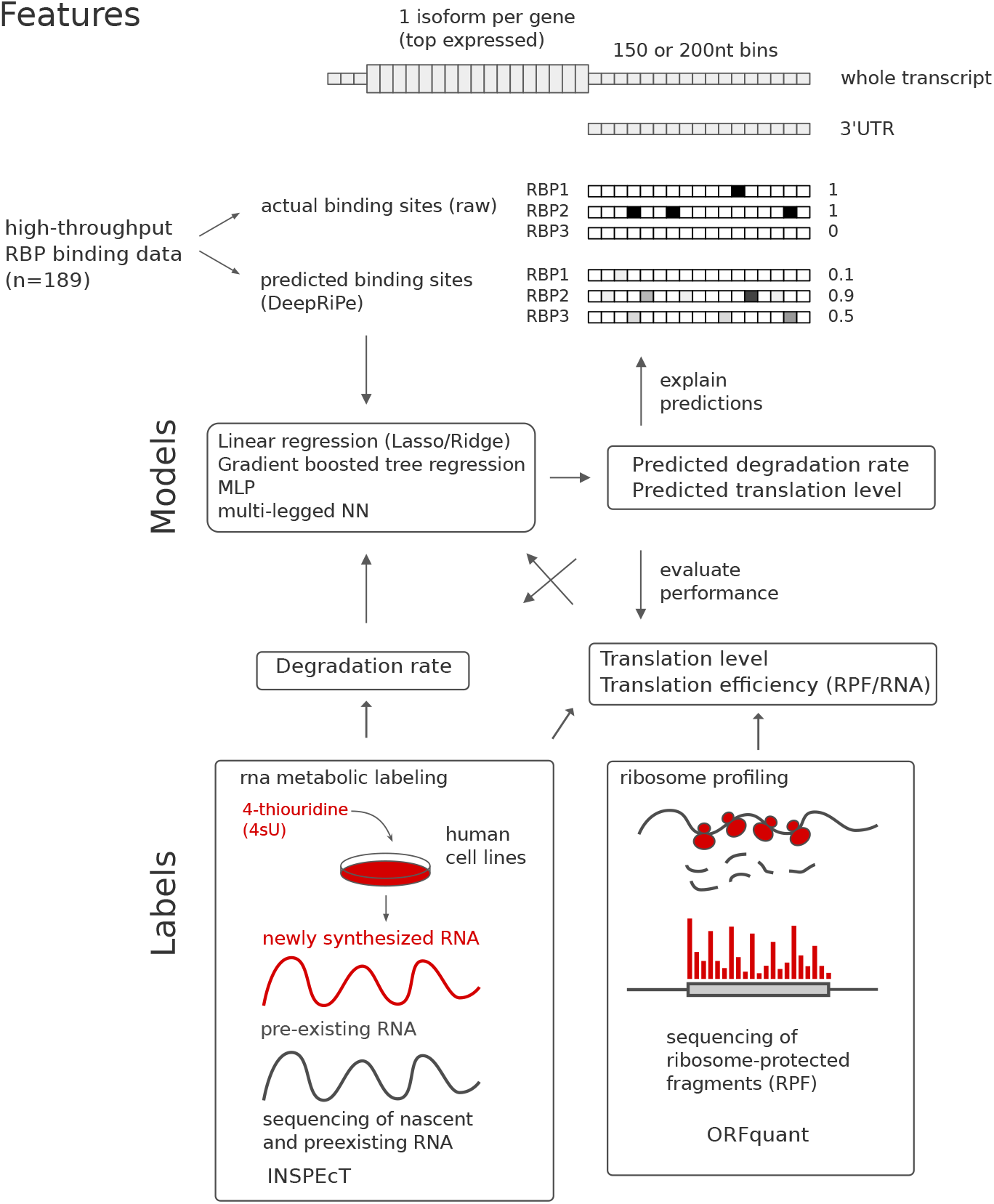
Overview of the prediction framework and dataset. *Top:* RBP features. For each gene, the top expressed transcript isoform is selected as the representative sequence. Experimental and predicted RBP binding sites derived from eCLIP and PAR-CLIP are summarized per transcript. *Bottom:* Prediction labels. *Middle:* Regression models used for prediction.

The output of the first step thus consists of aggregated raw CLIP and DeepRiPe features that will be the inputs for the following regression module, for either the whole transcript sequence or only the 3’UTR. Labels for the supervised training of the regressor are the RNA degradation rate estimates calculated from 4-thiouridine metabolic labeling experiments using INSPEcT [11]. We used the labeling data for four human cell lines: an exisiting HEK293 dataset [35], along with newly generated measurements in K562, HepG2, and HeLa cells. We derived gene-level estimates of degradation rates and assigned this value to the highest expressed isoform in each cell line. As labels for translation level, we utilized a value of “P sites per million” (ORFs pM) from ribosome profiling experiments [5]. In addition, we computed translation efficiency (TE) as a log-scale ratio between the ORFs pM and transcripts per million (TPM) derived from cytoplasmic RNA sequencing.

### 2.2. Learning a regressor

We evaluated several regression models, initially focusing on using training and test data from the same cell line (HEK293). We considered five models of different complexity levels: two linear regression models (lasso and ridge), gradient-boosted tree regressor (GBR), multi-layer perceptron (MLP), and two-legged neural network (the legs providing raw CLIP and DeepRiPe-predicted input separately into the network). Because translation is regulated by RNA-binding proteins at the 5’UTR, CDS, or 3’UTR, we use features for the whole transcript for interpretation of translation prediction. For predicting degradation rate, we focus on 3’UTR features only, as 3’UTR-binding RBPs are known to regulate RNA stability and informative features are easier to interpret.

We achieved a maximum performance of Pearson correlation between true and predicted values exceeding 0.75 for predicting translation level and 0.6 for degradation rate. Prediction performance dropped to 0.4 and 0.3, respectively, if only DeepRiPe features were used for prediction, but the best performance was achieved with the combination of both raw CLIP and DeepRiPe features. We observed small differences in the models’ performance (Figure 2) with cases where the simpler linear models and GBR outperformed neural networks, likely reflecting limitations of the dataset. As linear and tree-based models are easier to interpret and neural networks did not outperform them, we further focused on feature importance from ridge regression and GBR models to gain insight into which RBPs were important for predicting RNA stability and translation level.

**Figure 2:**
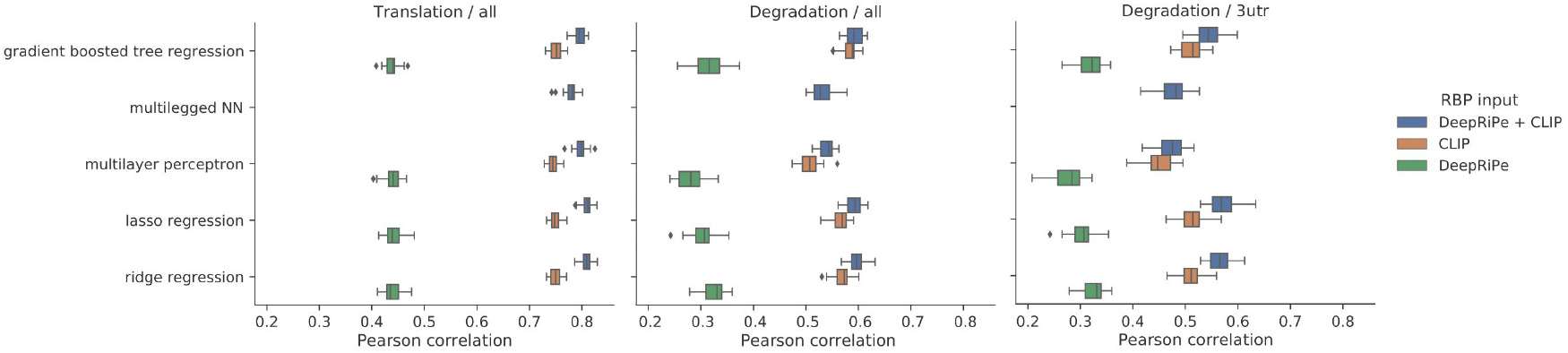
Predictive performance of regression models. X-axis: Average Pearson correlation coefficient between the true and predicted value of degradation rate or translation level in HEK293 cells for the five tested models. The right most plot is showing performance for models trained on 3’UTR region only, while for the the two other plots models are trained on the whole transcript. Color denotes the RBP feature set used in prediction: either DeepRiPe-predicted RBP sites (green), experimental RBP binding sites from eCLIP and PAR-CLIP (orange), or both (blue). Each model, label, and feature combination was run with five different random seeds.

### 2.3. Interpreting the results and the role of RBPs

We compared coefficients for ridge regression and feature importance from GBR prediction across several runs and identified the most important RNA-binding proteins to predict degradation rate and translation. The top 10 important features of GBR are shown in Figure 3 A, B 3.

**Figure 3:**
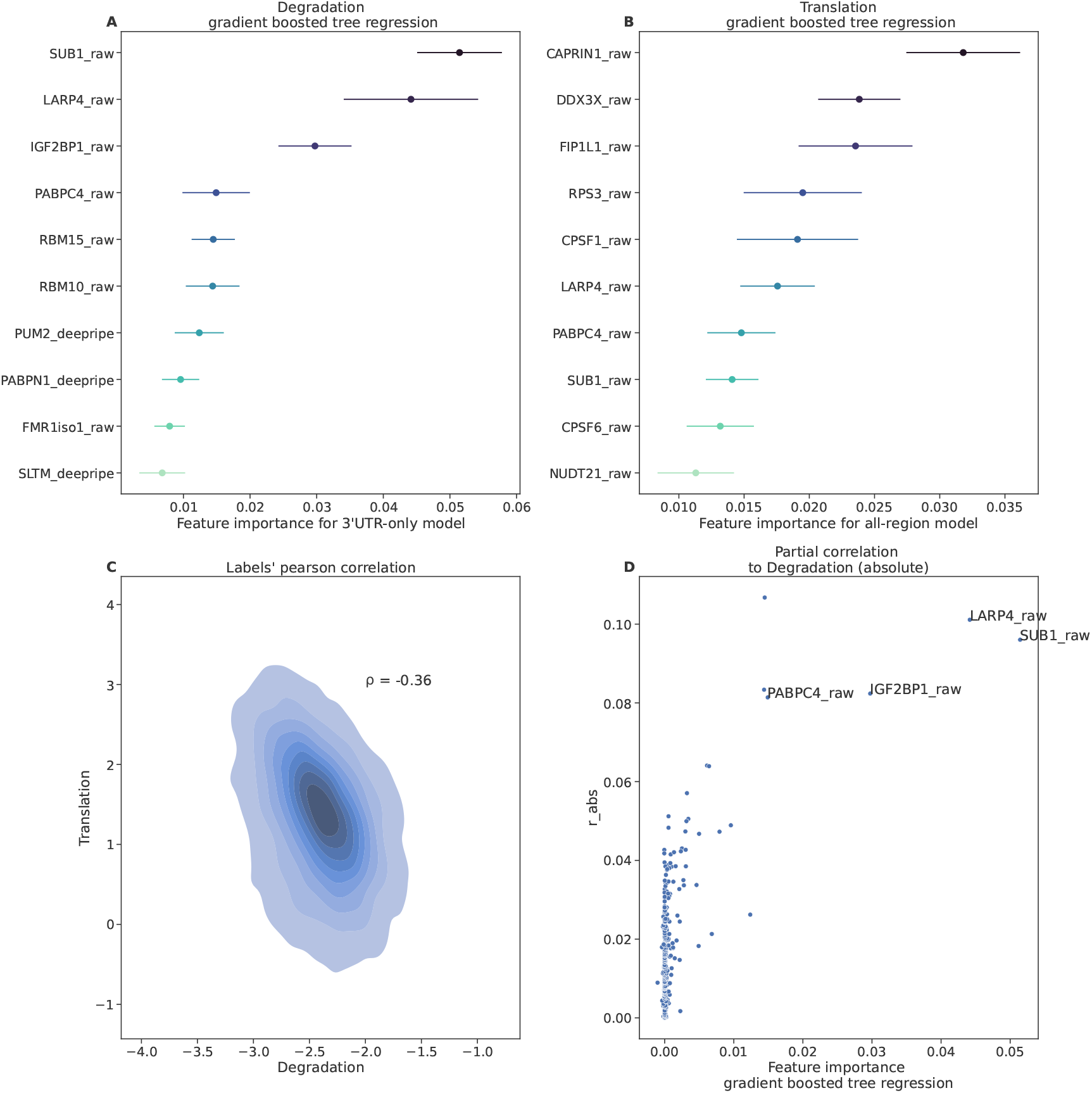
Top 10 important features for prediction of degradation with 3’UTR features (A) and translation with whole transcript features (B) with GBR. Error bars represent standard deviation from 5 runs initiated with different random seeds. (C) Correlation between degradation and transcript abundance. (D). Feature importance (GBR) against the partial correlation of each feature with degradation.

#### 2.3.1. RBPs for predicting degradation

GBR assigned high importance to both DeepRipe and raw CLIP features, whereas top ridge regression coefficients were highly dominated by raw features. Among the features with high importance, we find well known regulators of RNA stability: PUM2 destabilizes mRNA by recruiting CCR4-NOT deadenylase [16]; IGF2BP1 enhances mRNA stability by binding N6-methyladenosine (m6A) modified sites [24], and LARP4 stabilizes mRNA by binding to poly(A) tail and protecting from deadenylation [34]. Additionally, two more poly(A)-binding proteins, PABPC4 and PABPN1, also showed high importance, linking the degradation rate with the status of the poly(A) tail. Recent evidence shows that different transcripts have a strong bias towards different PABPs, which is reflected in their stability [38]. However, some important features do not have a known direct connection to cytoplasmic RNA stability, such as SUB1 and SLTM, nuclear transcription factors whose RNA-binding activity is only known from interactome studies ([2, 7]).

#### 2.3.2. RBPs for predicting translation

Similar to degradation, GBR gave high importance to RBPs known to be relevant for translation regulation. Thus, CAPRIN1 possibly regulates translation via condensates with FMRP1 [26]. Dead-box helicase DDX3X regulates translation as well [6]. Poly(A) binding proteins are again given high importance (LARP4, PABPN1). As for stability, there are again nuclear proteins among the imporant features, which are presumably not involved in translation but rather in transcription (SUB1) and cleavage and polyadenylation (FIP1L1, CPSF1).

#### 2.3.3. Degradation and translation are related

We wondered why our models would give some proteins high importance if they have no known role in degradation or translation. In addition, we noticed that three among the top ten important features selected by GBR for degradation prediction were among the top ten for predicting translation (LARP4, SUB1, PABPC1). To investigate this further, we looked at the relationship between those labels in our dataset. Indeed, the degradation rate is anticorrelated to translation level (Pearson = -0.49, Figure 3C 3). Moreover, both of these properties have a strong relationship to cytoplasmic RNA abundance (log transcript per million (TPM) value from the RNA sequencing of the cytoplasmic fraction) (Supplementary Figure A.7). We reasoned that the underlying correlation to RNA abundance may provide an explanation for shared RBP features that are highly important for prediction. In particular, partial correlation shows the correlation between two features while accounting for the effect of all other features. We thus computed the correlation of each RBP feature to the degradation rate while accounting for the effect of all other RBP features. The result is shown in Figure 3D 3. Indeed, the three most important features (LARP4, SUB1, PABPC1) also have the highest partial correlation to the degradation rate. All three of them are derived from raw CLIP data, and we reason that the underlying confounding RNA abundance, rather than their biological role in RNA stability, underlies their high importance for prediction. This also likely explains the distinctly lower performance of DeepRiPe versus CLIP-based features, as the computational predictions will be less confounded by RNA abundance.

#### 2.3.4. Cell line specific feature importance

Our newly generated RNA stability data allowed us to investigate if there were any cell line-specific RBPs regulating either RNA stability or translation. We trained models on datasets from different cell lines and focused specifically on DeepRiPe features, as raw RBP features may be confounded by the underlying transcript abundance. Comparing feature importance for K562, HepG2 and HeLa cell lines to HEK293, there is general agreement between feature importance for all of the cell lines (Figure 4). Thus, it is likely that there was no RBP with strong cell type-specific effects on RNA stability in our dataset.

**Figure 4:**
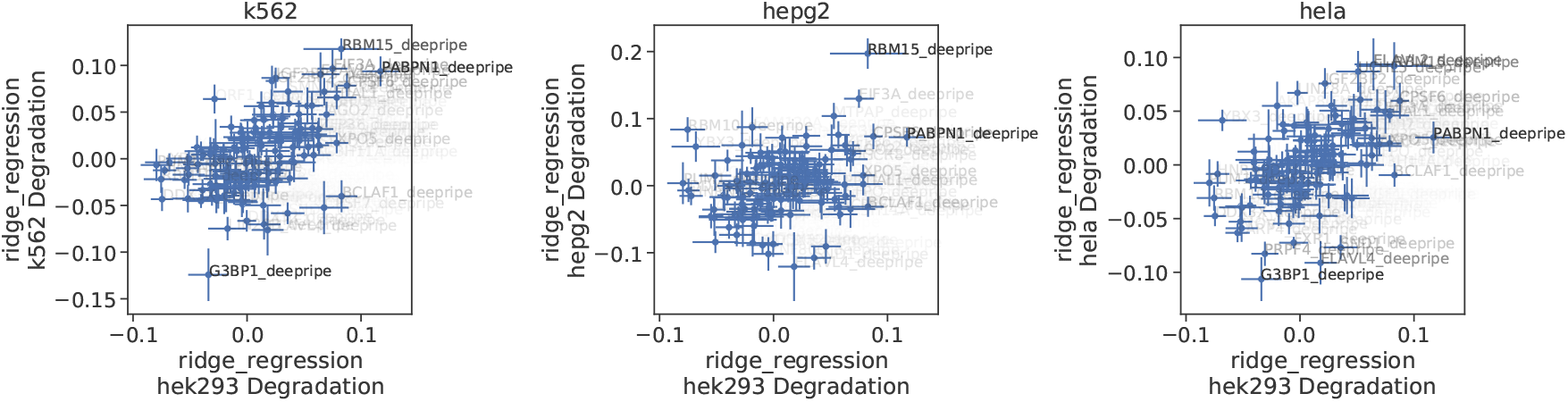
Comparison of feature importance (ridge regression coefficients) between models trained on HEK293 (X axis) and K562, HepG2 and HeLa, respectively (Y axis). Only DeepRiPe features are shown.

#### 2.3.5. Predicting of stability for unseen isoforms

Given that important RBP features were cell type agnostic, we asked if we could correctly predict the stability of isoforms that differ between cell lines. To this end, we used the model trained on the HEK293 dataset to predict RNA stability for those top expressed isoforms, which were present in the other three cell lines and did not occur in the HEK293 training set. The models achieved a Pearson correlation of approximately 0.3 (degradation rate) to 0.6 (translation level) (Supplementary Figure A.8). Examining correctly predicted isoforms (i.e., isoform among both the top N stable and top N predicted stable transcripts, similarly for unstable), we investigated individual predictions using Shapley values [31]. We asked if contributions of RBPs for individuals examples differed from the global feature importance and identified specific RBPs important for individual isoforms. We then used integrated gradients of a DeepRiPe model prediction [18] if a DeepRiPe feature was selected in the top 10 important for an isoform and the DeepRiPe prediction output was higher than 0.5. Figure 5 shows two examples where PUM2 is the most important feature for the prediction of degradation rates in a GBR model, and the DeepRiPe output is high for PUM2 in alternative parts of the transcript. The attribution map for this highly PUM2-positive bin clearly pinpoints an instance of the known PUM2 motif (UGUA(U/A)AUA) [17].

**Figure 5:**
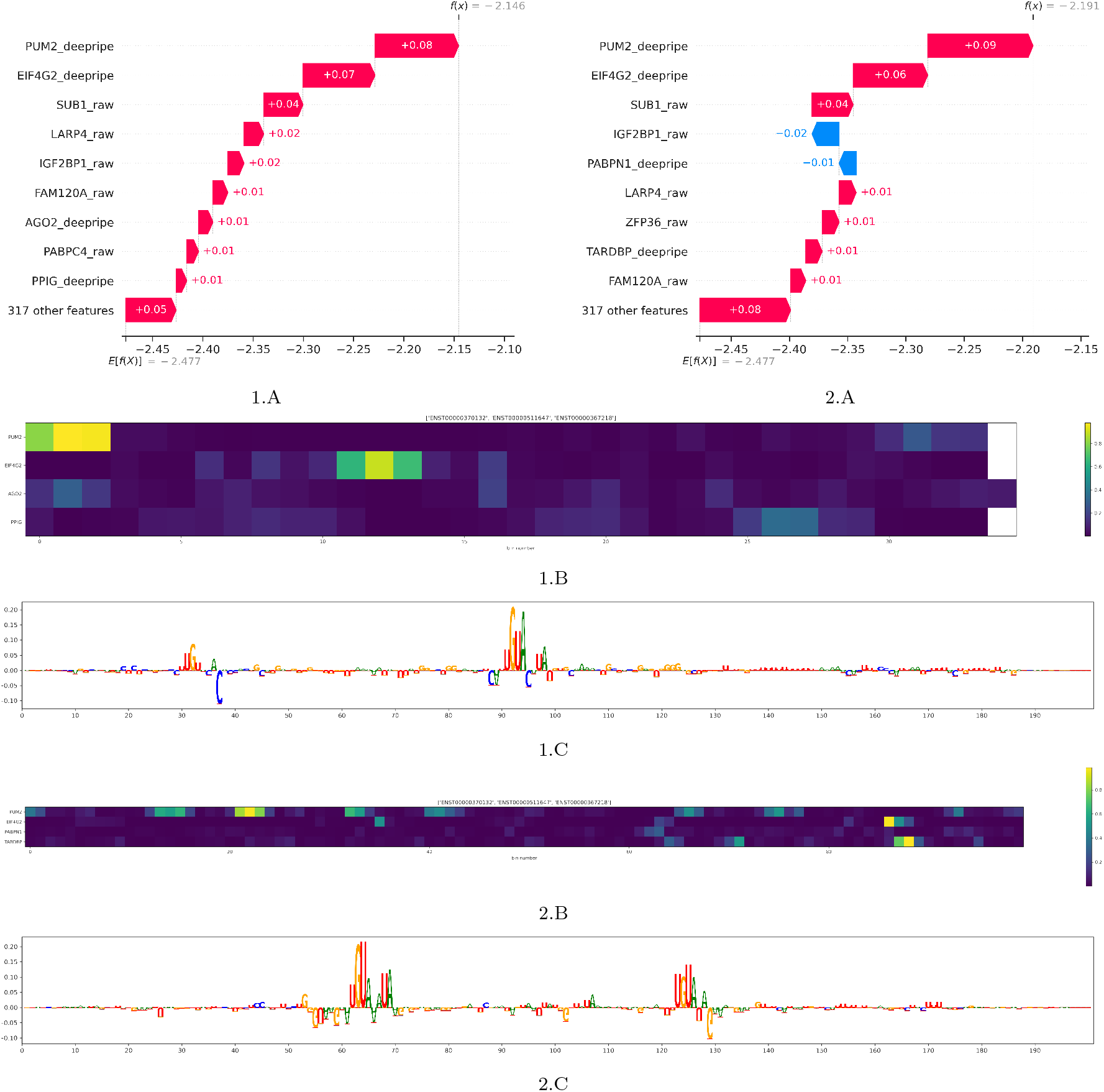
Feature importance analysis for two individual transcripts: ENST00000511647 and ENST00000370132. A: SHAP values. B: DeepRiPe prediction for 3UTR. C: Integrated gradients for PUM2 prediction for bin 50. 1.A: most important features for a HEK293 trained GBR model for degradation rates, and inference on transcript ENST00000511647 in the K562 cell line. 1.B: DeepRiPe binding prediction for overlapping sequence bins from transcript ENST00000511647, for selected RBPs. 1.C: attribution maps of DeepRiPe for PUM2 in the second sequence bin in transcript ENST00000511647. 2.A: most important features for a HEK293 trained GBR model for degradation rates, and inference on transcript ENST00000370132 in the K562 cell line. 2.B: binding prediction of DeepRiPe models for the first 100 overlapping sequence bins from transcript ENST00000370132, for selected RBPs. 2.C: attribution maps of DeepRiPe for class PUM2, at the sequence bin number 23 in transcript ENST00000370132 that is predicted strongly for PUM2 binding. Attribution maps 1.C and 2.C show known PUM2 motifs in the part of the transcript sequence where PUM2 binding is predicted by DeepRiPe.

#### 2.3.6. Predicting stability in controlled context

To evaluate the performance of stability predictors without confounding factors such as underlying alternative transcript abundance and codon composition, we turned to massively parallel reporter assay (MPRA) data [41]. In this assay, parts of 3’UTRs selected to contain AU-rich elements or other stability determinants were cloned into a lentiviral GFP reporter and transducted into the Jurkat cell line. RNA and DNA abundance was measured before and four hours after inducing transcriptional silencing of the reporter. The ratio of the RNA abundance between the two time points, normalized to DNA abundance, reflects the relative stability of the reporter carrying the specific 3’UTR insert. We predicted RBP binding sites on the inserts with DeepRiPe and used these features to predict reporter RNA stability with ridge regression and GBR. Here, we used available pre-trained DeepRipPe models for PAR-CLIP (as the size of the fragment exactly corresponded to the PAR-CLIP bin size but was too small for the eCLIP model). Despite the reduced amount of input features, we could predict stability with performance similar to that of prediction of degradation rate in transfer learning setting (Pearson correlation 0.3) (Figure 6A). Most important features selected by either model corresponded to ARE-binding proteins (ZFP36, ELAV2/3, ZC3H7B [29]) (Figure 6B).

**Figure 6:**
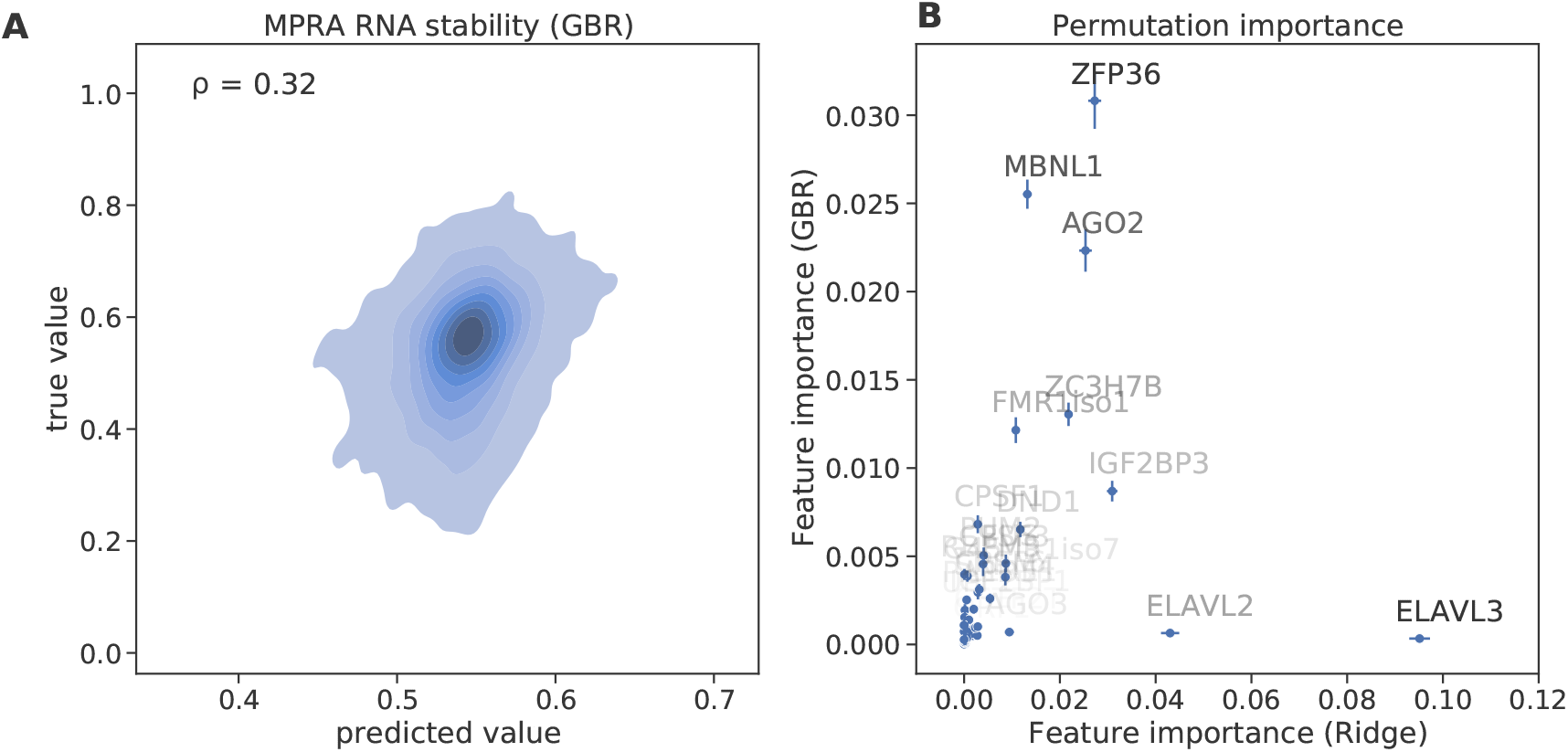
RNA stability prediction for massively parallel reporter assay (MPRA) [41]. (A) Correlation between true and predicted value for RNA stability for GBR prediction. (B) Comparison of top important features selected by GBR or Ridge models. Permutation importance is shown.

## 3. Discussion

### 3.1. Differences in feature importance between models

Given the data and task at hand, deep machine learning models did not exceed simpler choices, which are typically easier to interpret. We have compared highly important features of linear and GBR models. Whereas some features were specifically important for one model type, some had reproducibly high importance across models. Important features are expected to differ between linear regression and GBR, given the non-perfect performance of both models on the regression task. Additionally, the nature of the model plays a role. For example, SLBP protein has a very large coefficient in linear regressor for translation prediction (Supplemental Figure 1) because it exclusively binds histone mRNAs, which are highly translated, and is needed for their processing and translation [48]. However, only a few of the histone genes are expressed in a given cell type. Thus, this feature is not likely to be often used in the decision trees and has low importance in GBR. It is important to keep in mind that the absence of an RBP feature in the top feature list for a given model does not signify that this RBP is not important for the biological process, only that it was not helpful or redundant for a given model to obtain a good prediction.

### 3.2. Comparison of DeepRiPe and raw CLIP data

pWe included DeepRiPe features based on the assumption that these features may generalize better than raw CLIP data. First, in vivo binding data include false negatives due to low levels of transcript expression in the respective cell line. Additionally, raw CLIP data is not generally available across all cell lines and include false positives as well as non-specific binding sites, which will get a low prediction score from DeepRiPe if the RBP has considerable sequence specificity. While the performance using DeepRiPe features alone was at first seemingly worse than using raw CLIP data, this did not apply to the prediction of translation efficiency, for which translation levels are normalized to transcript abundance. This pointed to RNA abundance being a strongly confounding factor for CLIP-based prediction of stability and translation. Upon closer investigation, this was reflected in the set of most important features, which included repeatedly identified nuclear factors without known roles in stability or translation. While the methods to derive binding sites from CLIP data explicitly account for RNA abundance [46], there is apparent leakage for binding sites to be found on more highly expressed transcripts.

### 3.3. Relevance of using reporter assays

Despite their lower success, DeepRiPe-feature based models are more likely to correspond to the real ability to predict stablity and translation, and to identify biologically meaningful features. For instance, while of similarly limited performance, we were able to recover known AU-rich element binding proteins as most important features on the MPRA data. While not recapitulating the full regulatory sequence landscape, reporters have the advantage to focus on one aspect while keeping the rest constant, a useful characteristic to pinpoint important regulatory features in isolation.

### 3.4. Isoform-specific RNA stability

We derived degradation rates from single time-point metabolic labeling data, followed by modeling via INSPeCT [11]. This allows to perform experiments at lower costs, but comes at the price of having less precise estimates of RNA degradation rates. As it is inherently difficult to derive isoform-level stability measures from short-read data, we decided to use gene level estimates and assigned them to the most highly expressed isoform as a representative sequence.

The direct measurement of isoform-specific stability using long-read native RNA sequencing is therefore likely to be beneficial. We evaluated one such published dataset [33], and initial modelling results indeed exhibited a higher performance on these data. However, the half-lives reported by this study strongly diverge from those of other datasets we tested (for instance, ribosomal protein mRNAs have one of the shortest half-lives in this dataset). We therefore did not pursue this issue further, as it will have to be resolved by additional long-read derived stability data.

### 3.5. Limitations of the study

As outlined, the newly generated RNA stability data and assignment of the top expressed isoform sequence at a gene level is an approximation. We also decided to focus on two large datasets, the ENCODE and PAR-CLIP compendia, which have been processed in a consistent way. While additional RBP CLIP data is available and could be included as features, this still only covers a fraction of all RBPs. DeepRiPe performance varies across different RBPs [18], influenced both by experimental factors such as antibody quality, as well as the extent to which RBP target specification is accurately represented by the model. Thus, the advantages of a hierarchical approach, in which experimentally or computationally derived binding sites provide interpretable features to predict phenotypes, rather than prediction directly from the sequence, may be outweighed by the limitations of available data [1]. This situation warrants a deeper investigation of DNA and RNA foundation models [25, 9], which should represent a more complete picture of regulatory features and could then be fine-tuned for cell-type specific predictions of molecular phenotypes.

### 3.6. Conclusion

Machine learning models are now routinely used to predict gene regulatory mechanisms such as expression, chromatin states, or stability, either directly from sequence or, as we did here, from features derived from data or model predictions. With increasing complexity, model interpretation is deservedly gaining attention; caution is needed as the importance of a specific feature may not be directly related to biological function but to underlying confounding factors. Thus, computational development needs to be complemented by orthogonal datasets that allow for independent assessments, such as those from massively parallel reporter assays. Isolated sequence contexts and mutational analysis are helpful in discriminating important features as a step towards fully understanding the gene regulatory code in its native context.

## 4. Methods

### 4.1. Cell culture, metabolic labeling, RNA isolation

HeLa cells were cultured in low glucose DMEM (Thermo 31885023), HepG2 in high glucose DMEM (Thermo 41965039), and K562 in IMDM (Thermo 12440053), all supplemented with 10% FBS (Thermo 16000044), at 37°C with 5% CO_2_. Cells were passed every 2-3 days using TrypLE (Thermo A1217701) and PBS (Thermo 10010015) for washing. Total RNA was isolated using Trizol (Life Tech. 15596018) according to the manufacturer’s instructions Metabolic labeling was originally developed in [13]. 4-thiouridine (Enzo N-RP-2304-250) was added to the final concentration of 500*µ*M to the cell culture medium and cells were cultured for either 5 or 50 min. After that cells were washed with PBS and RNA was isolated with Trizol. Between 30 and 150 *µ*g RNA was used for subsequent steps. Biotinylation reaction was carried out in biotinylation buffer (10mM Tris-HCl pH 7.5 (Life Tech. 15567027), 1mM EDTA pH 8.0 (Life Tech. 15575020)) using 1*µ*l of 0.1 mg/ml Biotin-MTS stock solution (90066, Biotrend dissolved in DMF (Sigma D4551)) per *µ*g RNA. After rotating for 1.5h at room temperature, the RNA was extracted with chloroform using phase lock tubes (5 Prime 2302830) and re-precipitated using 1/10 vol. 5M NaCl (Life Tech. AM9760G) and 1 vol. isopropanol. To precipitate biotinylated RNA, it was denatured at 65°C for 10 min., placed immediately on ice, and subsequently rotated at room temperature with 100*µ*l streptavidin beads (Miltenyi 130-074-101). Columns (µMACS streptavidin kit 130-074-101) were equilibrated with washing buffer (100 mM Tris-Cl pH 7.5, 10 mM EDTA pH 8.0, 1 M NaCl, 0.1% Tween-20 (Sigma P2287)). The sample was applied to the column and washed with 4x 1ml preheated (65°C) and 4x room temperature washing buffer. RNA was eluted with 2x 100*µ*l 100 mM DTT (Sigma 43816) into 700 µl RNeasy MinElute RLT buffer (Qiagen 74204) and cleaned up using RNeasy MinElute kit according to the manufacturer’s instructions.

### 4.2. Cytoplasmic RNA

For HEK293, the same cytoplasmic RNA was used as in [35]. For other human cell lines, raw data for cytoplasmic RNA sequencing was downloaded from the ENCODE portal (https://www.encodeproject.org/) [32]. NCBI SRA IDs used were: SRR307929 for HepG2; SRR387661 for K562; SRR315334 for HeLa.

### 4.3. RNA sequencing

All RNA was processed with a RiboZero rRNA removal kit (Illumina MRZG12324). Subsequently, RNA sequencing libraries were prepared using NEXTflex RD qRNA-Seq Library Prep Kit (NOVA-5130-02D) according to the manufacturer’s instructions and sequenced on Illumina NextSeq 500.

### 4.4. Processing of RNA sequencing data

The pipeline for RNA sequencing read mapping is modified from [35] and can be found under https://github.com/slebedeva/rnaroids. The reference genome used for all human data is Gencode v19. Briefly, after UMI extraction with umi-tools [42], the reads are quality-trimmed using fastx-trimmer (http://hannonlab.cshl.edu/fastx_toolkit/). rRNA and ERCC reads are removed using bowtie [28]. Reads are mapped to the genome using STAR [12] and subsequently deduplicated with umi-tools. Deduplicated bam files were used to derive counts for degradation rate calculation. For the derivation of cytoplasmic TPM used to determine the highest expressed isoform as well as to calculate translation efficiency, kallisto [4] was used on deduplicated reads extracted after STAR alignment.

### 4.5. Derivation of degradation rates

Degradation rates were derived using the R package INSPEcT [11]. Gene-level degradation rates for all cell lines except HEK293 were derived using read counts from aligned bam files for total RNA and 50 min. labeled RNA. For HEK293, rates were averaged over 30, 45 and 60 min. labeling times. All degradation rates were log10-transformed.

### 4.6. Derivation of translation level and translation efficiency

ORFs pM values representing translation levels were derived directly from [5] (Supplemental Data 1). Translation efficiency (TE) was calculated by taking the log ratio of cytoplasmic RNA TPM and ORF pM. For calculating TE we selected transcripts with both TPM and ORFs pM exceeding 0.1. Both translation level and translation efficiency were log10-transformed. Cytoplasmic RNA TPM was obtained by running kallisto quantification on cytoplasmic RNA sequencing data from ENCODE project.

### 4.7. Derivation of RBP features

We used ENCODE eCLIP [47] and a collection of PAR-CLIP [36] RBP binding sites. We mapped RBP binding site centers to whole transcript or 3’UTR alone using the highest expressed isoform as a representative transcript per gene. The transcript sequence was separated into bins (200nt size for eCLIP and 150nt size for PAR-CLIP). To derive raw RBP features, we counted how many bins have an RBP center mapped to them per transcript and RBP. For DeepRipe features, we used pre-trained models from [18] to predict binding for each bin and RBP. Our model allows different ways of aggregating bin-level RBP features to transcript level (summarizing, taking maximum, or binarizing).

### 4.8. Partial correlation

Partial correlation between features and degradation label was derived with the python package pingoin [45].

### 4.9. Datasets

Depending on the selected configuration, our tool can predict either RNA degradation rate or translation level as the label, from different sets of input features. The first group of features, referred to as DeepRiPe RBP features, are the predicted binding scores from DeepRiPe for binding of each RBP, aggregated as the maximum score across all bins for each given transcript. The second group of features, if the binary score for binding of each RBP, according to the original CLIP data, aggregated as the maximum score for each transcript across all bins, eCLIP and PARCLIP data. We refer to this group as raw CLIP RBP features. Other features supported by our implementation include RBP expression and exon density. All the results presented in this manuscript are produced by taking DeepRiPe RBP features and raw CLIP RBP features jointly as input to the prediction models unless stated otherwise. Beyond the specified settings, our tool can be configured to choose from a variety of input data pre-processing options as well as model parameters.

### 4.10. Regression Models

We have benchmarked several of the most common regression machine learning models for predicting RNA degradation rate and translation rate from RBP binding information. We have experimented with training with k-fold cross-validation with multiple random seeds to assure the stability of the prediction. Furthermore, we have evaluated all models with fixed trainset and testsets split based on chromosome information to be reproducible and consistent with DeepRiPe’s training strategy. This section describes the specification of the regression models that are supported by our tool.

#### 4.10.1. Linear models

Linear models are our baseline for this prediction task, which include linear regression, lasso regression, and ridge regression. we use scikit-learn [39] implementation of these models in python together with minmax scaling of the input in the pipeline. We take the normalized coefficient of the trained model as an indicator of the learned feature importance to investigate the role of RBPs in the respective downstream task.

#### 4.10.2. Tree-based models

We use gradient boosting tree regressor (GBR) implemented by scikit-learn with min-max scaling of the input (and alternatively with XGBoost libraries). GBR is trained with 300 estimators with a maximum tree depth of 5 and early stopping of the training with a patience of 5.

#### 4.10.3. Neural Networks

We implement neural network models with various architectures to account for different ways of feeding input data into the model. Firstly, similar to our other model, we input concatenated and min-max-scaled DeepRiPe and raw CLIP features into a multi-layer perception (MLP) implemented using scikit-learn with min-max scaling. Second, we implement a two-legged neural network using Tensorflow, to feed each group of features separately, i.e. one leg for DeepRiPe-generated features and the other for raw CLIP features. Each leg has two hidden dense layers that are not shared with the other leg, and the legs are concatenated and followed by one last dense layer. The third implementation is a three-legged neural network, similar to the two-legged one, with an additional leg to take in exon density information. In all architectures, we use ReLu activation for neurons in the hidden layers and we train for a maximum of 50 epochs with early-stopping enabled with a patience of 5. For MLP we use the Adam optimizer and for our multi-legged networks, we use Nadam for optimization.

## 5. Code access

All code is freely available at https://github.com/sepidehsaran/rbp-based-stability-regressor.

## 6. Data access

All raw and processed sequencing data generated in this study will be made publicly available through the NCBI Gene Expression Omnibus (GEO).

## Acknowledgements

The authors are grateful to Neelanjan Mukherjee for planning out metabolic labeling data collection and providing the analysis pipeline. Thanks to Sophia Bauch for planning out HeLa metabolic labeling experiments. We acknowledge Mahsa Ghanbari for contributing to machine learning design and discussions.

## 9. Author contributions

U.O., S.L., and S.S. designed the study. A.H. performed metabolic labeling experiments and sequencing. S.L. performed data preprocessing, exploratory analysis, and model interpretation. S.S. performed modeling and wrote the code. S.L., S.S., and U.O. wrote the paper.

## Appendix A. Supplementary Materials

**Figure A.7:**
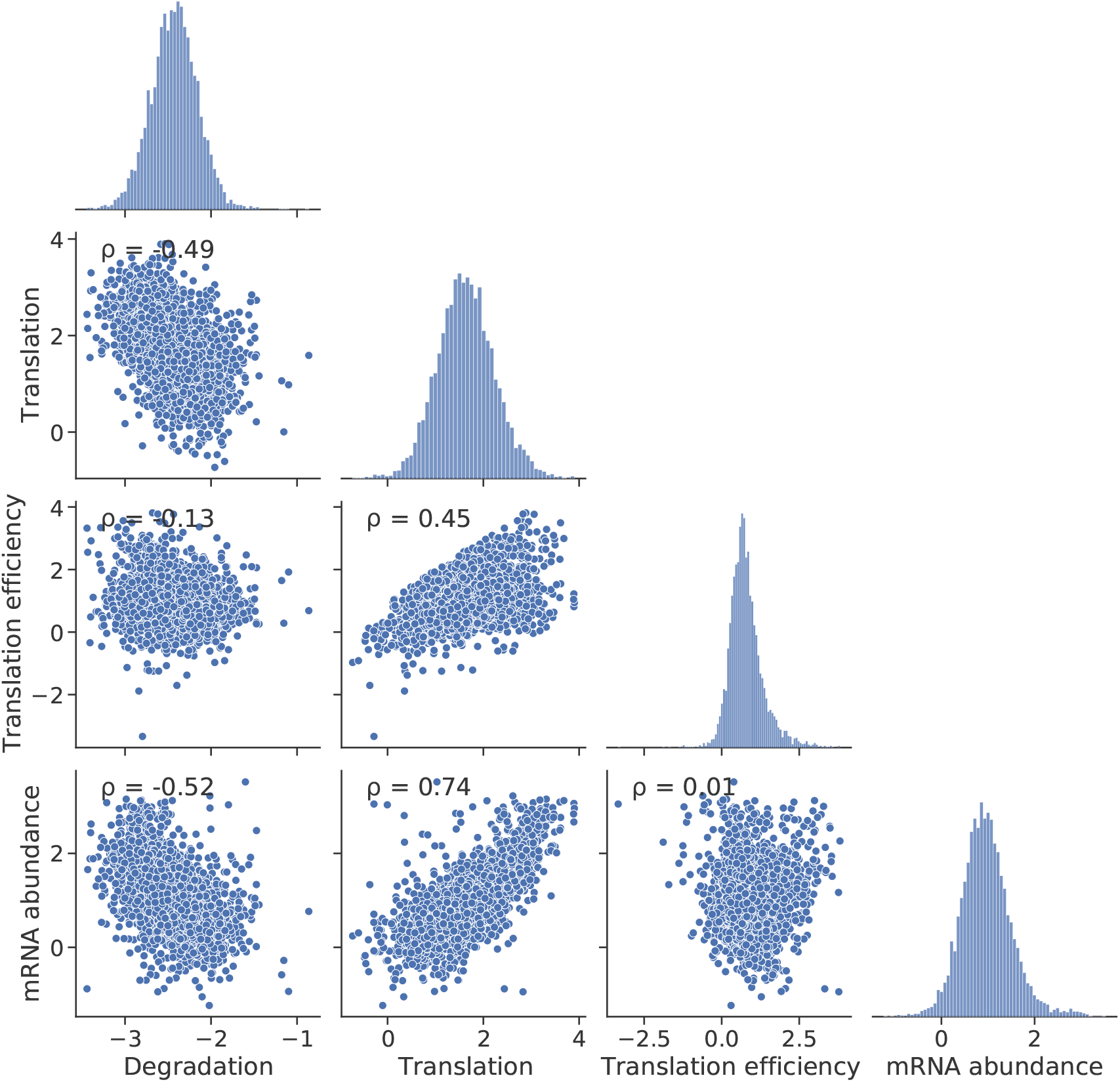
Correlation between prediction labels: degradation, translation and translation efficiency (TE) as well as RNA abundance (represented by log10(transcripts per million (TPM)) derived from kallisto[4]). Numbers denote Pearson correlation coefficient.

**Figure A.8:**
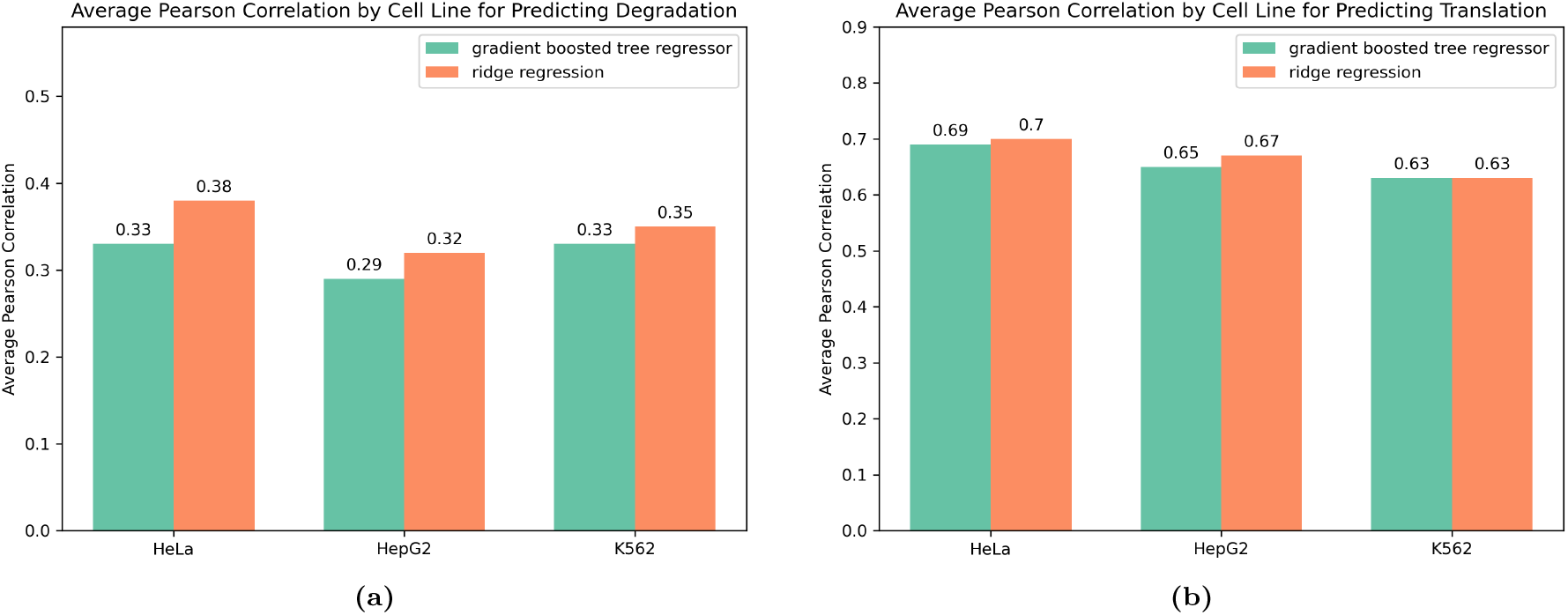
Performance of the two best model types (i.e. gradient boosted tree and ridge regression) on the transfer learning task. All models were trained on HEK293 dataset and tested on isoforms specific for the denoted cell line. (a) average pearson correlation for predicting Degradation rate over five random seeds. (b) average pearson correlation for predicting Translation rate over five random seeds.

## References

[1] V. Agarwal and D. Kelley. The genetic and biochemical determinants of mRNA degradation rates in mammals. URL https://www.biorxiv.org/content/10.1101/2022.03.18.484474v1. Pages: 2022.03.18.484474 Section: New Results.

[2] A. G. Baltz, M. Munschauer, B. Schwanhäusser, A. Vasile, Y. Murakawa, M. Schueler, N. Youngs, D. Penfold-Brown, K. Drew, M. Milek, E. Wyler, R. Bonneau, M. Selbach, C. Dieterich, and M. Landthaler. The mRNA-bound proteome and its global occupancy profile on protein-coding transcripts. 46(5):674–690. ISSN 1097-2765. doi: 10.1016/j.molcel.2012.05.021. URL https://www.sciencedirect.com/science/article/pii/S1097276512004376.

[3] K. W. Brannan, I. A. Chaim, R. J. Marina, B. A. Yee, E. R. Kofman, D. A. Lorenz, P. Jagannatha, K. D. Dong, A. A. Madrigal, J. G. Underwood, and G. W. Yeo. Robust single-cell discovery of RNA targets of RNA-binding proteins and ribosomes. Nature Methods, 18(5):507–519, May 2021. ISSN 1548-7105. doi: 10.1038/s41592-021-01128-0. URL https://www.nature.com/articles/s41592-021-01128-0. Number: 5 Publisher: Nature Publishing Group.

[4] N. L. Bray, H. Pimentel, P. Melsted, and L. Pachter. Near-optimal probabilistic RNA-seq quantification. 34(5):525–527. ISSN 1546-1696. doi: 10.1038/nbt.3519. URL https://www.nature.com/articles/nbt.3519. Number: 5 Publisher: Nature Publishing Group.

[5] L. Calviello, A. Hirsekorn, and U. Ohler. Quantification of translation uncovers the functions of the alternative transcriptome. 27(8):717–725,. ISSN 1545-9985. doi: 10.1038/s41594-020-0450-4. URL https://www.nature.com/articles/s41594-020-0450-4. Number: 8 Publisher: Nature Publishing Group.

[6] L. Calviello, S. Venkataramanan, K. J. Rogowski, E. Wyler, K. Wilkins, M. Tejura, B. Thai, J. Krol, W. Filipowicz, M. Landthaler, and S. Floor. DDX3 depletion represses translation of mRNAs with complex 5 UTRs. 49(9):5336–5350,. ISSN 0305-1048. doi: 10.1093/nar/gkab287. URL 10.1093/nar/gkab287.

[7] A. Castello, B. Fischer, K. Eichelbaum, R. Horos, B. M. Beckmann, C. Strein, N. E. Davey, D. T. Humphreys, T. Preiss, L. M. Steinmetz, J. Krijgsveld, and M. W. Hentze. Insights into RNA biology from an atlas of mammalian mRNA-binding proteins. 149(6):1393–1406. ISSN 0092-8674, 1097-4172. doi: 10.1016/j.cell.2012.04.031. URL https://www.cell.com/cell/abstract/S0092-8674(12)00576-4. Publisher: Elsevier.

[8] J. Cheng, K. C. Maier, Avsec, P. Rus, and J. Gagneur. Cis-regulatory elements explain most of the mRNA stability variation across genes in yeast. 23(11):1648–1659. ISSN 1355-8382, 1469-9001. doi: 10.1261/rna.062224.117. URL http://rnajournal.cshlp.org/content/23/11/1648. Company: Cold Spring Harbor Laboratory Press Distributor: Cold Spring Harbor Laboratory Press Institution: Cold Spring Harbor Laboratory Press Label: Cold Spring Harbor Laboratory Press Publisher: Cold Spring Harbor Lab.

[9] Y. Chu, D. Yu, Y. Li, K. Huang, Y. Shen, L. Cong, J. Zhang, and M. Wang. A 5 UTR language model for decoding untranslated regions of mRNA and function predictions. 6(4): 449–460. ISSN 2522-5839. doi: 10.1038/s42256-024-00823-9. URL https://www.nature.com/articles/s42256-024-00823-9. Publisher: Nature Publishing Group.

[10] C. C. Friedel and L. Dölken. Metabolic tagging and purification of nascent RNA : implications for transcriptomics. 5(11):1271–1278. doi: 10.1039/B911233B. URL https://pubs.rsc.org/en/content/articlelanding/2009/mb/b911233b. Publisher: Royal Society of Chemistry.

[11] S. de Pretis, T. Kress, M. J. Morelli, G. E. M. Melloni, L. Riva, B. Amati, and M. Pelizzola. INSPEcT: a computational tool to infer mRNA synthesis, processing and degradation dynamics from RNA- and 4su-seq time course experiments. 31(17):2829–2835. ISSN 1367-4803. doi: 10.1093/bioinformatics/btv288. URL 10.1093/bioinformatics/btv288.

[12] A. Dobin, C. A. Davis, F. Schlesinger, J. Drenkow, C. Zaleski, S. Jha, P. Batut, M. Chaisson, and T. R. Gingeras. STAR: ultrafast universal RNA-seq aligner. 29(1):15–21. ISSN 1367-4803. doi: 10.1093/bioinformatics/bts635. URL 10.1093/bioinformatics/bts635.

[13] L. Dölken. High resolution gene expression profiling of RNA synthesis, processing, and decay by metabolic labeling of newly transcribed RNA using 4-thiouridine. 1064:91–100. ISSN 1940-6029. doi: 10.1007/978-1-62703-601-66.

[14] K.N. D’Orazio and R. Green. Ribosome states signal RNA quality control. 81(7):1372–1383. ISSN 1097-2765. doi: 10.1016/j.molcel.2021.02.022. URL https://www.ncbi.nlm.nih.gov/pmc/articles/PMC8041214/.

[15] T. J. Eisen, S. W. Eichhorn, A. O. Subtelny, K. S. Lin, S. E. McGeary, S. Gupta, and D. P. Bartel. The dynamics of cytoplasmic mRNA metabolism. 77(4):786–799.e10. ISSN 1097-2765. doi: 10.1016/j.molcel.2019.12.005. URL https://www.cell.com/molecular-cell/abstract/S1097-2765(19)30896-2. Publisher: Elsevier.

[16] J. V. Etten, T. L. Schagat, J. Hrit, C. A. Weidmann, J. Brumbaugh, J. J. Coon, and A. C. Goldstrohm. Human pumilio proteins recruit multiple deadenylases to efficiently repress messenger RNAs *. 287(43):36370–36383. ISSN 0021-9258, 1083-351X. doi: 10.1074/jbc.M112.373522. URL https://www.jbc.org/article/S0021-9258(20)62611-4/abstract. Publisher: Elsevier.

[17] A. Galgano, M. Forrer, L. Jaskiewicz, A. Kanitz, M. Zavolan, and A. P. Gerber. Comparative analysis of mRNA targets for human PUF-family proteins suggests extensive interaction with the miRNA regulatory system. 3(9):e3164. ISSN 1932-6203. doi: 10.1371/journal.pone.0003164. URL https://journals.plos.org/plosone/article?id=10.1371/journal.pone.0003164. Publisher: Public Library of Science.

[18] M. Ghanbari and U. Ohler. Deep neural networks for interpreting RNA-binding protein target preferences. 30(2):214–226. ISSN 1088-9051. doi: 10.1101/gr.247494.118. URL https://www.ncbi.nlm.nih.gov/pmc/articles/PMC7050519/.

[19] A. Guvenek, J. Shin, L. De Filippis, D. Zheng, W. Wang, Z. P. Pang, and B. Tian. Neuronal cells display distinct stability controls of alternative polyadenylation mRNA isoforms, long non-coding RNAs, and mitochondrial RNAs. 13. ISSN 1664-8021. URL https://www.frontiersin.org/articles/10.3389/fgene.2022.840369.

[20] M. Hafner, M. Katsantoni, T. Köster, J. Marks, J. Mukherjee, D. Staiger, J. Ule, and M. Zavolan. CLIP and complementary methods. 1(1):1–23. ISSN 2662-8449. doi: 10.1038/s43586-021-00018-1. URL https://www.nature.com/articles/s43586-021-00018-1. Number: 1 Publisher: Nature Publishing Group.

[21] E. Hartenian and B. A. Glaunsinger. Feedback to the central dogma: cytoplasmic mrna decay and transcription are interdependent processes. 54(4):385–398. ISSN 1040-9238. doi: 10.1080/10409238.2019.1679083. URL https://www.ncbi.nlm.nih.gov/pmc/articles/PMC6871655/.

[22] V. A. Herzog, B. Reichholf, T. Neumann, P. Rescheneder, P. Bhat, T. R. Burkard, W. Wlotzka, A. von Haeseler, J. Zuber, and S. L. Ameres. Thiol-linked alkylation of RNA to assess expression dynamics. Nature Methods, 14(12):1198–1204, Dec 2017. ISSN 1548-7105. doi: 10.1038/nmeth.4435. URL https://www.nature.com/articles/nmeth.4435. Number: 12 Publisher: Nature Publishing Group.

[23] M. Horlacher, G. Cantini, J. Hesse, P. Schinke, N. Goedert, S. Londhe, L. Moyon, and A. Marsico. A systematic benchmark of machine learning methods for protein-RNA interaction prediction. 24(5):bbad307. ISSN 1477-4054. doi: 10.1093/bib/bbad307.

[24] H. Huang, H. Weng, W. Sun, X. Qin, H. Shi, H. Wu, B. S. Zhao, A. Mesquita, C. Liu, C. L. Yuan, Y.-C. Hu, S. Hüttelmaier, J. R. Skibbe, R. Su, X. Deng, L. Dong, M. Sun, C. Li, S. Nachtergaele, Y. Wang, C. Hu, K. Ferchen, K. D. Greis, X. Jiang, M. Wei, L. Qu, J.-L. Guan, C. He, J. Yang, and J. Chen. Recognition of RNA n6-methyladenosine by IGF2bp proteins enhances mRNA stability and translation. 20(3):285–295. ISSN 1476-4679. doi: 10.1038/s41556-018-0045-z. URL https://www.nature.com/articles/s41556-018-0045-z. Number: 3 Publisher: Nature Publishing Group.

[25] A. Karollus, J. Hingerl, D. Gankin, M. Grosshauser, K. Klemon, and J. Gagneur. Species-aware DNA language models capture regulatory elements and their evolution. 25:83. doi: 10.1186/s13059-024-03221-x. URL https://pmc.ncbi.nlm.nih.gov/articles/PMC10985990/.

[26] T. H. Kim, B. Tsang, R. M. Vernon, N. Sonenberg, L. E. Kay, and J. D. Forman-Kay. Phospho-dependent phase separation of FMRP and CAPRIN1 recapitulates regulation of translation and deadenylation. 365(6455):825–829. doi: 10.1126/science.aax4240. URL https://www.science.org/doi/10.1126/science.aax4240. Publisher: American Association for the Advancement of Science.

[27] W. S. Lai, R. M. Arvola, A. C. Goldstrohm, and P. J. Blackshear. Inhibiting transcription in cultured metazoan cells with actinomycin d to monitor mRNA turnover. 155:77–87. ISSN 1046-2023. doi: 10.1016/j.ymeth.2019.01.003. URL https://www.ncbi.nlm.nih.gov/pmc/articles/PMC6392460/.

[28] B. Langmead, C. Trapnell, M. Pop, and S. L. Salzberg. Ultrafast and memory-efficient alignment of short DNA sequences to the human genome. 10(3):R25. ISSN 1474-760X. doi: 10.1186/gb-2009-10-3-r25. URL 10.1186/gb-2009-10-3-r25.

[29] K. Leppek and G. Stoecklin. An optimized streptavidin-binding RNA aptamer for purification of ribonucleoprotein complexes identifies novel ARE-binding proteins. 42(2):e13. ISSN 1362-4962. doi: 10.1093/nar/gkt956.

[30] A. Lugowski, B. Nicholson, and O. S. Rissland. Determining mRNA half-lives on a transcriptome-wide scale. 137:90–98. ISSN 1095-9130. doi: 10.1016/j.ymeth.2017.12.006.

[31] S. M. Lundberg and S.-I. Lee. A unified approach to interpreting model predictions. In Advances in Neural Information Processing Systems, volume 30. Curran Associates, Inc. URL https://papers.nips.cc/paper_files/paper/2017/hash/8a20a8621978632d76c43dfd28b67767-Abstract.html.

[32] Y. Luo, B. C. Hitz, I. Gabdank, J. A. Hilton, M. S. Kagda, B. Lam, Z. Myers, P. Sud, J. Jou, K. Lin, U. K. Baymuradov, K. Graham, C. Litton, S. R. Miyasato, J. S. Strattan, O. Jolanki, J.-W. Lee, F. Y. Tanaka, P. Adenekan, E. O’Neill, and J. M. Cherry. New developments on the encyclopedia of DNA elements (ENCODE) data portal. 48:D882–D889. ISSN 0305-1048. doi: 10.1093/nar/gkz1062. URL 10.1093/nar/gkz1062.

[33] K. C. Maier, S. Gressel, P. Cramer, and B. Schwalb. Native molecule sequencing by nano-ID reveals synthesis and stability of RNA isoforms. 30(9):1332–1344. ISSN 1549-5469. doi: 10.1101/gr.257857.119.

[34] S. Mattijssen, G. Kozlov, B. D. Fonseca, K. Gehring, and R. J. Maraia. LARP1 and LARP4: up close with PABP for mRNA 3’ poly(a) protection and stabilization. 18(2):259–274. ISSN 1555-8584. doi: 10.1080/15476286.2020.1868753.

[35] N. Mukherjee, L. Calviello, A. Hirsekorn, S. de Pretis, M. Pelizzola, and U. Ohler. Integrative classification of human coding and noncoding genes through RNA metabolism profiles. 24 (1):86–96,. ISSN 1545-9985. doi: 10.1038/nsmb.3325. URL https://www.nature.com/articles/nsmb.3325. Number: 1 Publisher: Nature Publishing Group.

[36] N. Mukherjee, H.-H. Wessels, S. Lebedeva, M. Sajek, M. Ghanbari, A. Garzia, A. Munteanu, D. Yusuf, T. Farazi, J. I. Hoell, K. M. Akat, A. Akalin, T. Tuschl, and U. Ohler. Deciphering human ribonucleoprotein regulatory networks. 47(2):570–581,. ISSN 1362-4962. doi: 10.1093/nar/gky1185.

[37] M. Müller-McNicoll, O. Rossbach, J. Hui, and J. Medenbach. Auto-regulatory feedback by RNA-binding proteins. 11(10):930–939. ISSN 1759-4685. doi: 10.1093/jmcb/mjz043. URL 10.1093/jmcb/mjz043.

[38] A. L. Nicholson-Shaw, E. R. Kofman, G. W. Yeo, and A. E. Pasquinelli. Nuclear and cytoplasmic poly(a) binding proteins (PABPs) favor distinct transcripts and isoforms. 50(8): 4685–4702. ISSN 1362-4962. doi: 10.1093/nar/gkac263.

[39] F. Pedregosa, G. Varoquaux, A. Gramfort, V. Michel, B. Thirion, O. Grisel, M. Blondel, P. Prettenhofer, R. Weiss, V. Dubourg, J. Vanderplas, A. Passos, D. Cournapeau, M. Brucher, M. Perrot, and E. Duchesnay. Scikit-learn: Machine learning in Python. Journal of Machine Learning Research, 12:2825–2830, 2011.

[40] M. Schmid and T. H. Jensen. Controlling nuclear RNA levels. 19(8):518–529. ISSN 1471-0064. doi: 10.1038/s41576-018-0013-2. URL https://www.nature.com/articles/s41576-018-0013-2. Number: 8 Publisher: Nature Publishing Group.

[41] D. A. Siegel, O. Le Tonqueze, A. Biton, N. Zaitlen, and D. J. Erle. Massively parallel analysis of human 3 UTRs reveals that AU-rich element length and registration predict mRNA destabilization. 12(1):jkab404. ISSN 2160-1836. doi: 10.1093/g3journal/jkab404. URL https://www.ncbi.nlm.nih.gov/pmc/articles/PMC8728028/.

[42] T. S. Smith, A. Heger, and I. Sudbery. UMI-tools: Modelling sequencing errors in unique molecular identifiers to improve quantification accuracy. page gr.209601.116. ISSN 1088-9051, 1549-5469. doi: 10.1101/gr.209601.116. URL https://genome.cshlp.org/content/early/2017/01/18/gr.209601.116. Company: Cold Spring Harbor Laboratory Press Distributor: Cold Spring Harbor Laboratory Press Institution: Cold Spring Harbor Laboratory Press Label: Cold Spring Harbor Laboratory Press Publisher: Cold Spring Harbor Lab.

[43] F. X. R. Sutandy, S. Ebersberger, L. Huang, A. Busch, M. Bach, H.-S. Kang, J. Fallmann, D. Maticzka, R. Backofen, P. F. Stadler, K. Zarnack, M. Sattler, S. Legewie, and J. König. In vitro iCLIP-based modeling uncovers how the splicing factor u2af2 relies on regulation by cofactors. 28(5):699–713. ISSN 1088-9051, 1549-5469. doi: 10.1101/gr.229757.117. URL https://genome.cshlp.org/content/28/5/699. Company: Cold Spring Harbor Laboratory Press Distributor: Cold Spring Harbor Laboratory Press Institution: Cold Spring Harbor Laboratory Press Label: Cold Spring Harbor Laboratory Press Publisher: Cold Spring Harbor Lab.

[44] Y. Uchida, T. Chiba, R. Kurimoto, and H. Asahara. Post-transcriptional regulation of inflammation by RNA-binding proteins via cis-elements of mRNAs. 166(5):375–382. ISSN 0021-924X. doi: 10.1093/jb/mvz067. URL https://www.ncbi.nlm.nih.gov/pmc/articles/PMC6804641/.

[45] R. Vallat. Pingouin: statistics in python. 3(31):1026. ISSN 2475-9066. doi: 10.21105/joss.01026. URL https://joss.theoj.org/papers/10.21105/joss.01026.

[46] E. L. Van Nostrand, P. Freese, G. A. Pratt, X. Wang, X. Wei, R. Xiao, S. M. Blue, J.-Y. Chen, N. A. L. Cody, D. Dominguez, S. Olson, B. Sundararaman, L. Zhan, C. Bazile, L. P. B. Bouvrette, J. Bergalet, M. O. Duff, K. E. Garcia, C. Gelboin-Burkhart, M. Hochman, N. J. Lambert, H. Li, M. P. McGurk, T. B. Nguyen, T. Palden, I. Rabano, S. Sathe, R. Stanton, Su, R. Wang, B. A. Yee, B. Zhou, A. L. Louie, S. Aigner, X.-D. Fu, E. Lécuyer, C. B. Burge, R. Graveley, and G. W. Yeo. A large-scale binding and functional map of human RNA-binding proteins. 583(7818):711–719,. ISSN 1476-4687. doi: 10.1038/s41586-020-2077-3. URL https://www.nature.com/articles/s41586-020-2077-3. Number: 7818 Publisher: Nature Publishing Group.

[47] E. L. Van Nostrand, G. A. Pratt, A. A. Shishkin, C. Gelboin-Burkhart, M. Y. Fang, B. Sundararaman, S. M. Blue, T. B. Nguyen, C. Surka, K. Elkins, R. Stanton, F. Rigo, M. Guttman, and G. W. Yeo. Robust transcriptome-wide discovery of RNA-binding protein binding sites with enhanced CLIP (eCLIP). 13(6):508–514,. ISSN 1548-7105. doi: 10.1038/nmeth.3810. URL https://www.nature.com/articles/nmeth.3810. Number: 6 Publisher: Nature Publishing Group.

[48] M. L. Whitfield, H. Kaygun, J. A. Erkmann, W. H. D. Townley-Tilson, Z. Dominski, and W. F. Marzluff. SLBP is associated with histone mRNA on polyribosomes as a component of the histone mRNP. 32(16):4833–4842. ISSN 0305-1048. doi: 10.1093/nar/gkh798. URL https://www.ncbi.nlm.nih.gov/pmc/articles/PMC519100/.

